## Supplementary information for "Structure of cyanobacterial photosystem I complexed with Cytochrome *c*_6_ and Ferredoxin at 1.97 Å resolution"

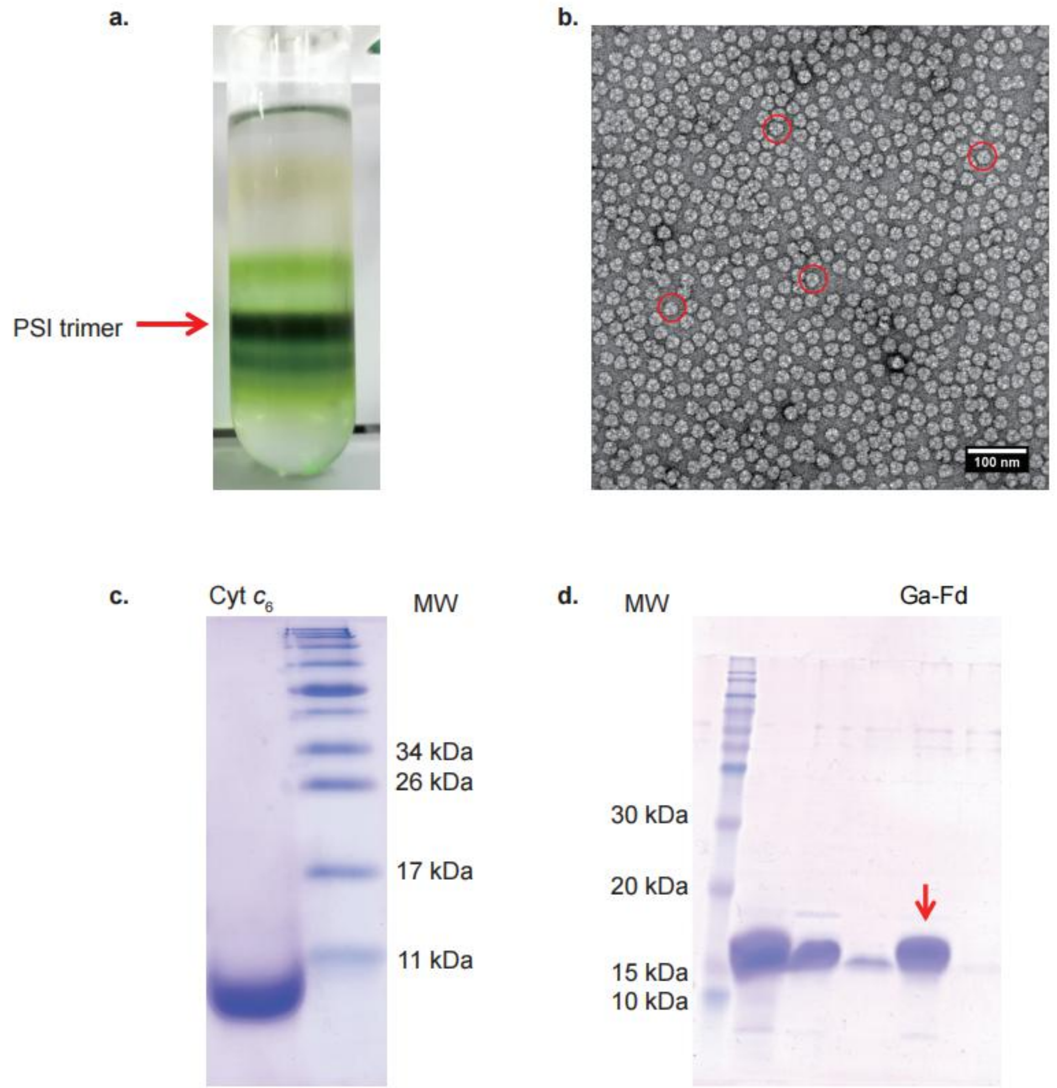


**Supplementary Figure 1. Purification of the PSI trimer, Ga-Fd and Cyt $c_6$. a.** Sucrose density gradient centrifugation of PSI after Ni-NTA affinity chromatography. The band corresponding to PSI trimers is highlighted by a red arrow. **b.** Micrograph of a negatively stained PSI trimer preparation. Representative PSI trimer particles with easy to discern protomers in the top-view are circled in red. **c.** SDS-PAGE of purified native Cyt $c_6$. **d.** SDS-PAGE of purified Ga-substituted Ferredoxin (Ga-Fd). The Coomassie blue stained band corresponding to the final purification step is marked by a red arrow.

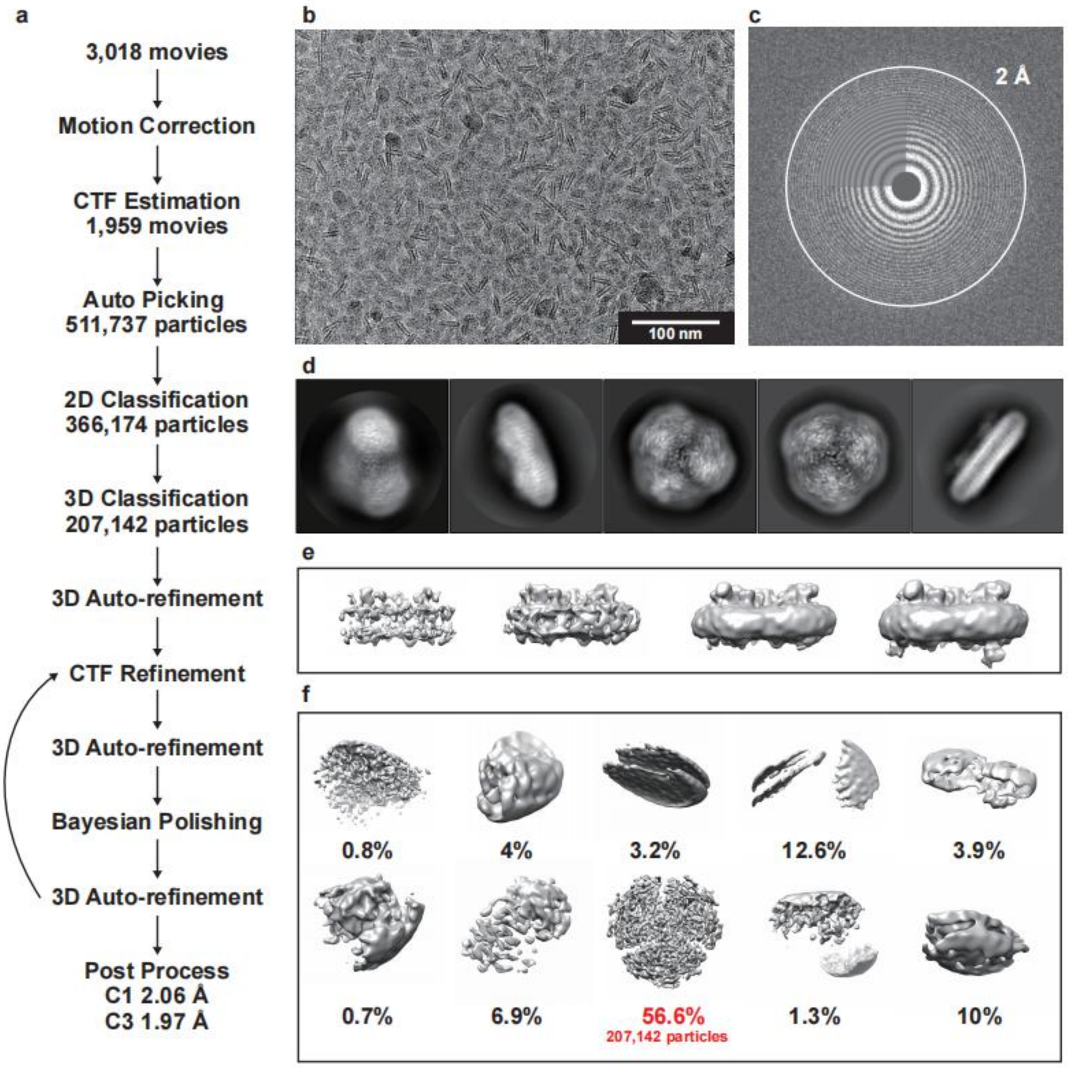


**Supplementary Figure 2. Workflow and results of single particle cryo image processing. a.** The workflow of data processing was performed in RELION 3.1. **b.** A representative micrograph and **c.** its Fourier transform with Thon rings extending to 2.0 Å resolution. **d.** 2D classes selected for further 3D classification. **e.** Initial *de novo* model based on auto-picked particles rendered at four different thresholds. **f.** The results of 3D classification. Particles in the best class (colored in red) were used for 3D refinement.

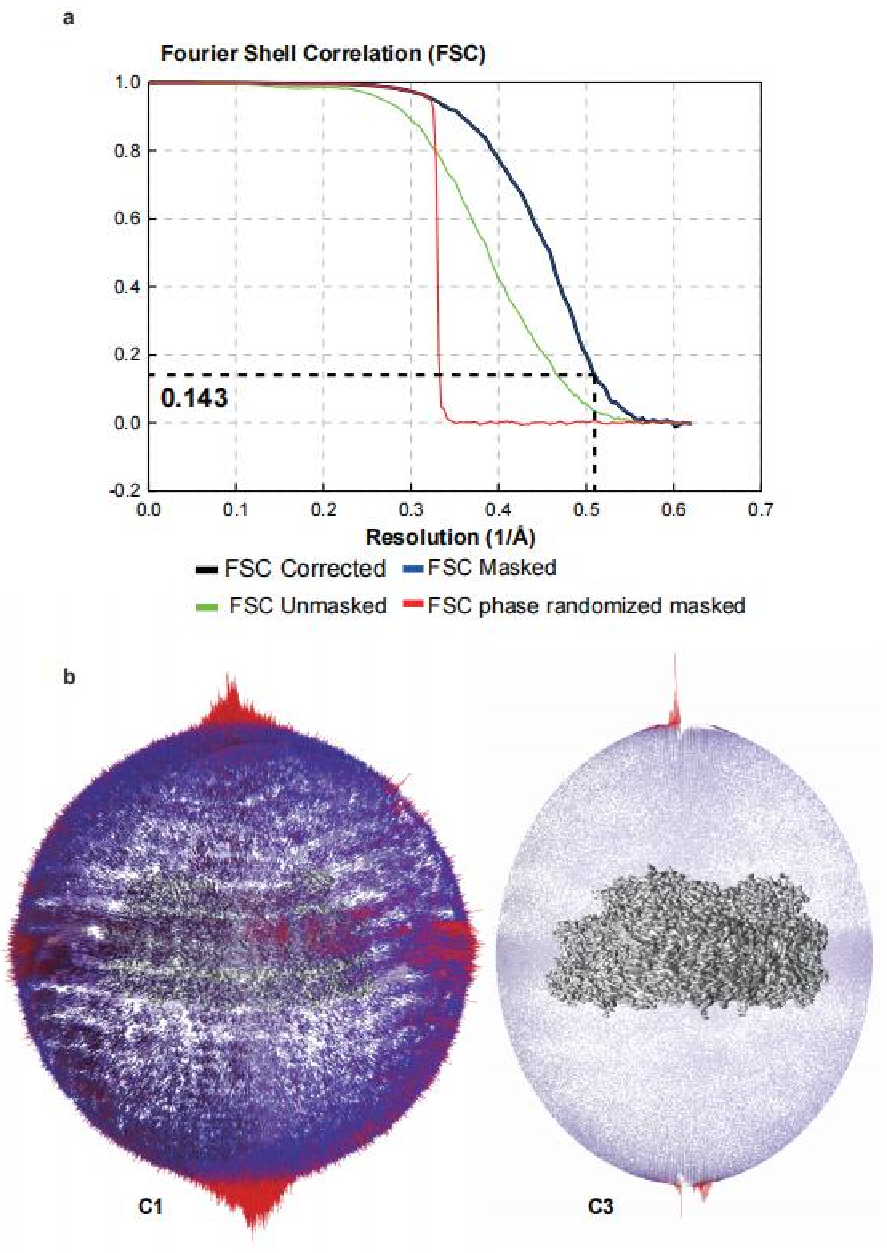


**Supplementary Figure 3. The FSC curves and Euler angle distribution. a.** The Gold Standard Fourier Shell Correlation (FSC) curves as implemented in Relion 3.1 for the post-processing results of the final map in C3 symmetry. The overall resolution of the masked density map was determined as 1.97 Å at 0.143 Gold Standard FSC cut-off. **b.** The Euler angle distribution of the final cryo-EM maps in C1 (left) and C3 (right) symmetry.

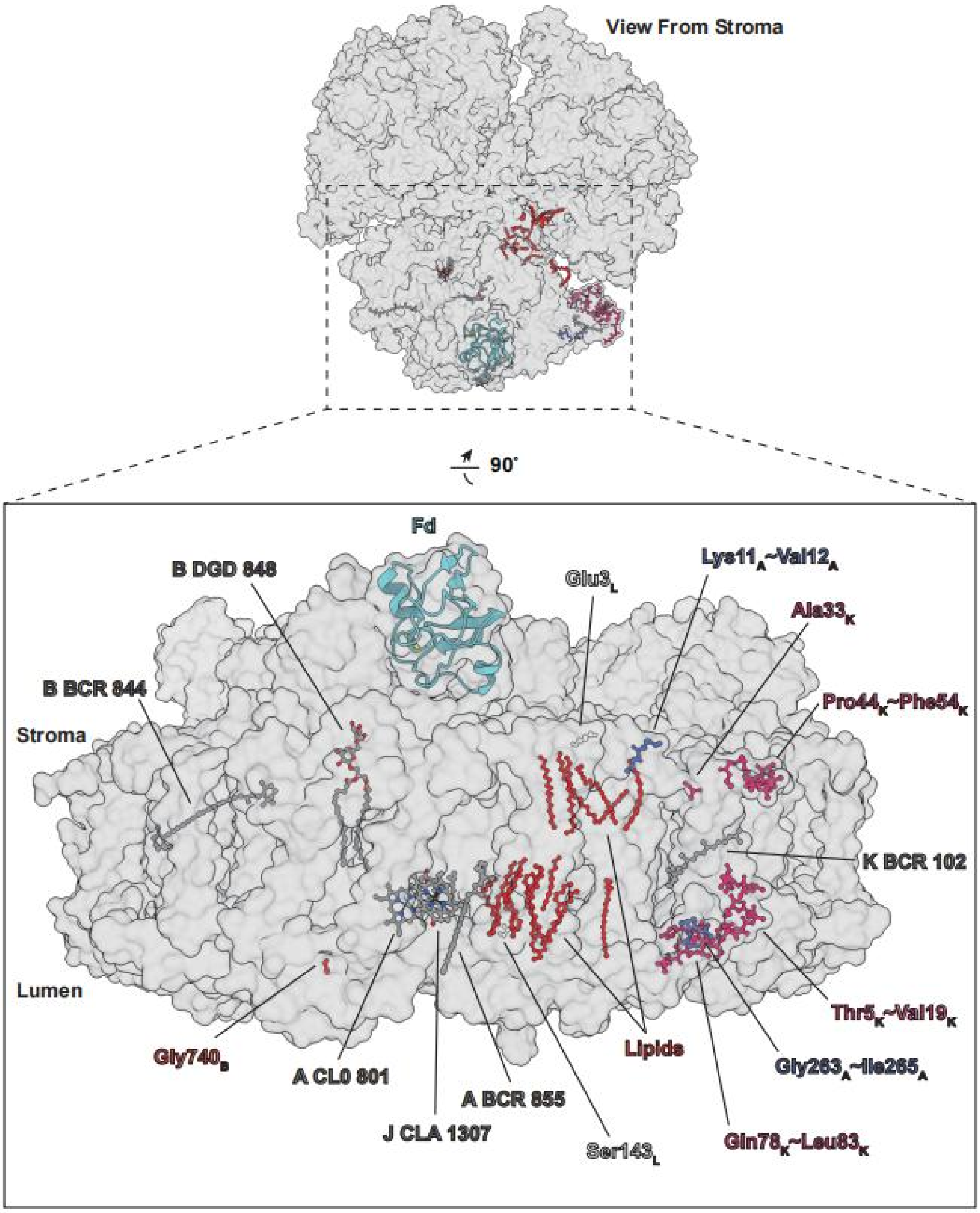


**Supplementary Figure 4. Summary of all newly added and modified residues/ligands as presented for one PSI protomer.** Additionally, bound Fd is presented in ribbon cartoon and transparent surface. Amino acid residues are colored according to subunits as in Figure 1. Ligands are colored by elements and lipids are shown in red.

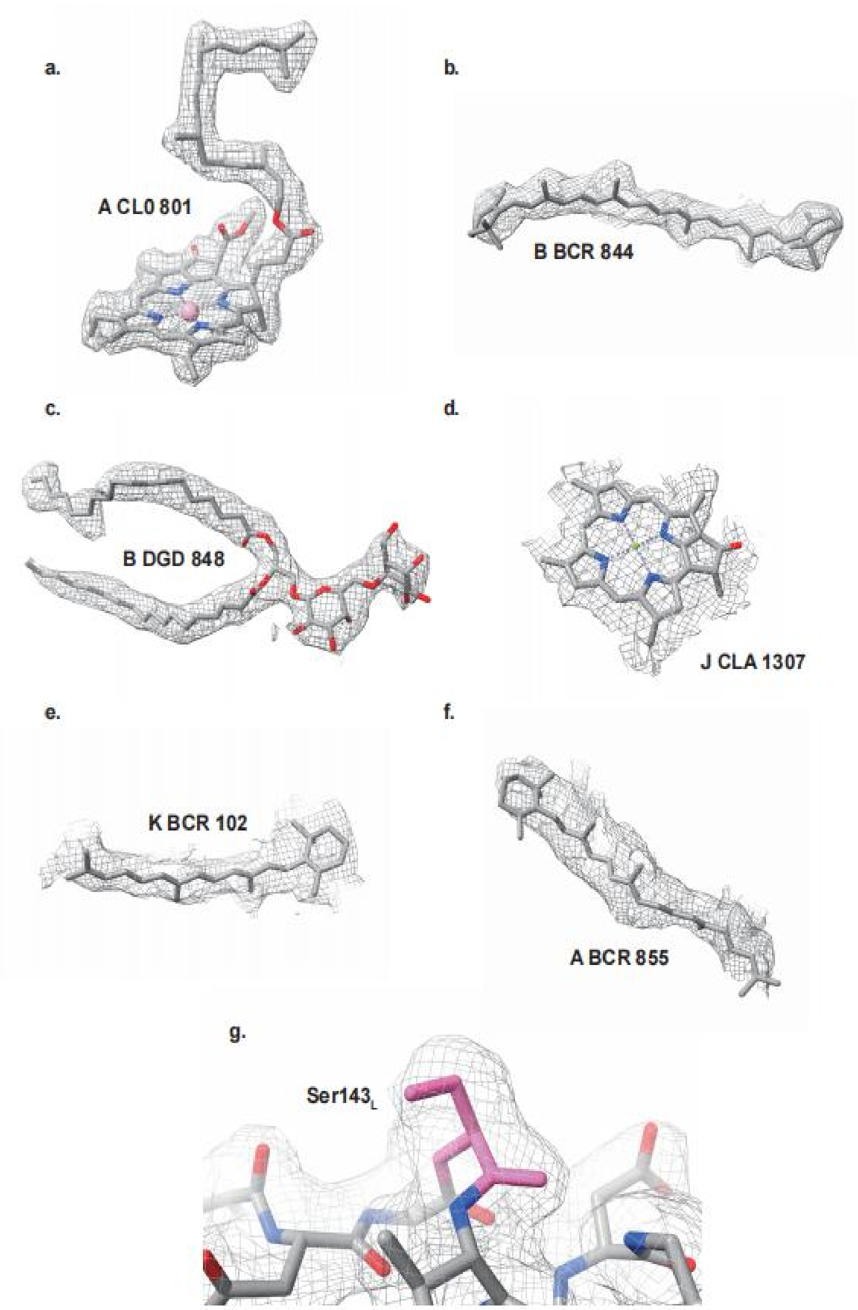


**Supplementary Figure 5. Representative additions and modifications to the starting model 1JB0 and their corresponding density maps. a.** One chlorophyll of the P700 chlorophyll pair is assigned as the chlorophyll *a'* isomer (PDB Ligands category CL0). **b.** β-carotene B844 was completely built with all atoms. **c.** Previously assigned as 1,2-distearoyl-monogalactosyl-diglyceride (MGDG, PDB Ligands category LMG) B848 was modified to digalactosyldiacylglycerol (DGDG, PDB Ligands category DGD) based on the newly detected extra density in the headgroup region. **d.** Newly assigned chlorophyll J1307 and corresponding density. **e.** and **f.** are newly assigned, partly built β-carotene K102 and A855, respectively. **g.** The 143th amino acid residue (colored in pink) in PSI subunit PsaL was modified from leucine to serine based on the Uniprot sequencing result and corresponding density map.

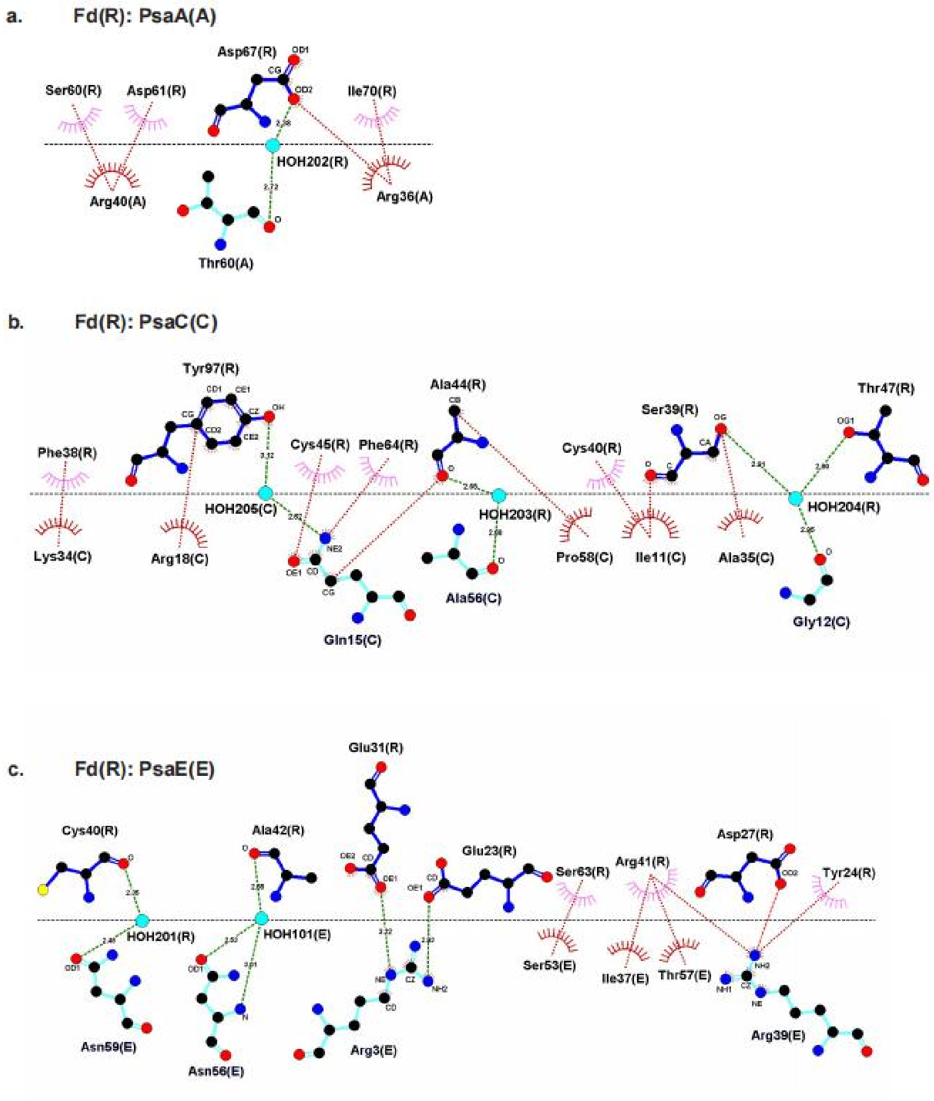


**Supplementary Figure 6. Protein-protein interface diagrams of a. Fd:PsaA, b. Fd:PsaC and c. Fd:PsaE derived from DIMPLOT of the LigPlot+ suite.** Green dashed lines represent potential hydrogen bonds. Red dashed lines show potential non-bonded contacts such as hydrophobic or cation-π interactions.

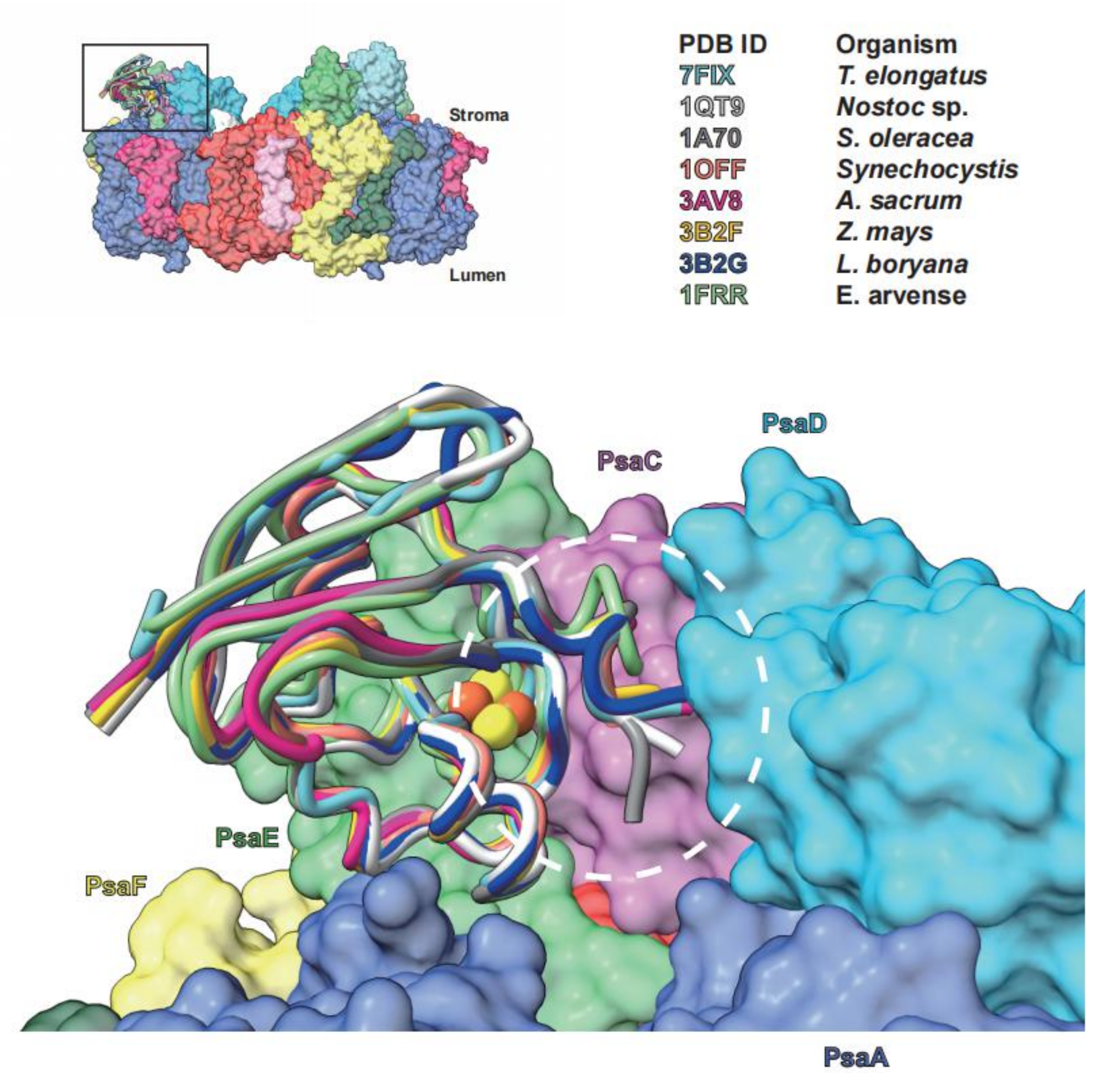


**Supplementary Figure 7. Superposition of Ferredoxin structures from diverse sources illustrates the structural flexibility of the C-terminus (white dashed circle).**

| **PSI:Fd complex (EMDB: 31605) (PDB: 7FIX) (EMPIAR: 10928)** | |
|---|---|
| **Data collection and processing** | |
| Microscope | CRYO ARM 300 (300 kV) (JEOL) |
| Camera | Gatan K3 |
| Defocus range (μm) | 0.5-1.5 |
| Electron dose (frame/total) ($e^-$/$Å^2$) | 1 / 48 |
| Pixel size (Å) | 0.806 |
| Exposure time (s) | 3 |
| Micrograph movies (initial/final) | 3,018 / 1,959 |
| Magnification | ×60,000 |
| Particles (initial/final) | 367,967 / 207,142 |
| Symmetry imposed | C1 / C3 |
| FSC threshold | 0.143 |
| Map resolution (Å) | 2.06 (C1) / 1.97 (C3) |
| **Refinement** | |
| Initial model used (PDB code) | 1JB0 (PSI), 5AUI (Fd) |
| R.M.S.D. (bonds (Å)/angles (°)) | 0.01 / 1.059 |
| Clashscore | 5.68 |
| Ramachandran favored (%) | 97.92 |
| Ramachandran allowed (%) | 2.06 |
| Ramachandran outliers (%) | 0 |
| Rotamer outliers (%) | 0 |
| Cβ outliers (%) | 0 |
| MolProbity score | 1.33 |

**Supplementary Table 1. Cryo-EM data collection, refinement and validation statistics.**

| | 1st measurement | 2nd measurement | 3rd measurement | Mean ± S.E.M. |
|---|---|---|---|---|
| $K_d$ (nM) | 703 | 994 | 577 | 758.0 ± 123.5 |
| Δ*G* (kcal/mol) | -8.4 | -8.2 | -8.5 | -8.4 ± 0.1 |
| Δ*H* (kcal/mol) | 1.0 | 1.1 | 1.2 | 1.1 ± 0.1 |
| -TΔS (kcal/mol) | -9.4 | -9.3 | -9.7 | -9.5 ± 0.1 |
| n | 2.9 | 3.2 | 2.5 | 2.9 ± 0.2 |

**Supplementary Table 2. Thermodynamic parameters of the interaction between Ga-Fd and PSI.** Three independent ITC experiments were performed at 25°C. Each set of thermodynamic parameters obtained by fitting normalized ΔH in the binding isotherm are shown without fit errors. Mean and S.E.M. values are also presented, indicating the averaged value and the standard error of the mean, respectively. Kd: dissociation constant; ΔG: change in Gibbs free energy; ΔH: enthalpy change; ΔS: change in entropy; n: binding stoichiometry.

| Author | Publish year | Protein | Organism | Mutation site (Corresponding residue in 7FIX) | Method | Potential influence |
|---|---|---|---|---|---|---|
| Jonathan Hanley et al[1] | 1996 | PSI PsaD | Synechocystis 6803 | H97 (H95), K106 (K104), R111 (R109) | Flash-absorption spectroscopy | Fd binding affinity with PSI |
| Nicolas Fischer et al[2] | 1998 | PSI PsaC | C. reinhardtii | **K35 (K34)** | Flash-absorption spectroscopy<br>Electron paramagnetic resonance spectroscopy | Fd binding affinity with PSI |
| Tetsuyuki Akashi et al[3] | 1999 | Fd | Z. mays | E93 | Cyclic voltammetry | Fd redox potential |
| Tetsuyuki Akashi et al[3] | 1999 | Fd | Z. mays | D66, **D67** | Affinity chromatography | Fd interaction with FNR and SiR |
| Nicolas Fischer et al[4] | 1999 | PSI PsaC | C. reinhardtii | D9 (D8), **I12 (I11)**, T15 (T14), **Q16 (Q15)** | Flash-absorption spectroscopy<br>Electron paramagnetic resonance spectroscopy | Fd binding affinity with PSI |
| Patrick Barth et al[5] | 2000 | PSI PsaE | Synechocystis 6803 | **R39** | Flash-absorption spectroscopy | Fd binding affinity with PSI |
| Bernard Lagoutte et al[6] | 2001 | PSI PsaD | Synechocystis 6803 | R111 (R109) | NADP+ Photo-reduction Assay<br>Flash-absorption spectroscopy | Fd binding affinity with PSI |
| Herve Bottin et al[7] | 2001 | PSI PsaD | Synechocystis 6803 | D100 (D98), E105 (E103), E109 (K107) | Flash-absorption spectroscopy | Fd binding affinity with PSI |
| Pierre Setif et al[8] | 2002 | PSI PsaC | T. elongatus | **K34**, G36 | Flash-absorption spectroscopy | Fd binding affinity with PSI |
| Hisako Kubota-Kawai et al[9] | 2018 | Fd | T. elongatus | **Y24, E31, D61, D67,** E71, Y81, E93, **Y97** | Flash-absorption spectroscopy | Fd binding affinity with PSI |

**Supplementary Table 3. A summary of site directed mutation studies on the interaction of PSI and Fd.** The residues colored in red were identified as being involved in the interaction with PSI in this study.
